# ICoRD: Iterative correlation-based ROI detection method for the extraction of neural signals in calcium imaging

**DOI:** 10.1101/2021.12.16.473055

**Authors:** Seongtak Kang, Jiho Park, Kyungsoo Kim, Sung-Ho Lim, Samhwan Kim, Joon Ho Choi, Jong-Cheol Rah, Ji-Woong Choi

**Affiliations:** Daegu Gyeongbuk Institute of Science and Technology (DGIST); University of California, San Francisco (UCSF); KEPCO Engineering & Construction Company (KEPCO E&;C); Korea Brain Research Institute (KBRI)

**Keywords:** signal-to-noise power ratio, region of interest, calcium imaging, denoising

## Abstract

*In vivo* calcium imaging is a standard neuroimaging technique that allows selective observation of target neuronal activities. In calcium imaging, neuron activation signals provide key information for the investigation of neural circuits. For efficient extraction of the calcium signals of neurons, selective detection of the region of interest (ROI) pixels corresponding to the active subcellular region of the target neuron is essential. However, current ROI detection methods for calcium imaging data exhibit a relatively low signal extraction performance from neurons with a low signal-to-noise power ratio (SNR). This is problematic because a low SNR is unavoidable in many biological experiments. Therefore, we propose an iterative correlation-based ROI detection (ICoRD) method that robustly extracts the calcium signal of the target neuron from a calcium imaging series with severe noise. ICoRD extracts calcium signals closer to the ground-truth calcium signal than the conventional method from simulated calcium imaging data in all low SNR ranges. Additionally, this study confirmed that ICoRD robustly extracts activation signals against noise, even within *in vivo* environments. ICoRD showed reliable detection from neurons with a low SNR and sparse activation, which were not detected by conventional methods. ICoRD will facilitate our understanding of neural circuit activity by providing significantly improved ROI detection in noisy images.

## 1. Introduction

Intracellular calcium concentration remains low in the resting state and increases abruptly upon neuronal activity [1-4]. Therefore, calcium imaging is a standard neuroimaging method for measuring neuronal activity in the living brain [5-8]. For the precise extraction of information on activity, a selective grouping of pixels corresponding to the intracellular calcium change of a target structure, such as the cell body or dendrite, is required. The group of pixels has been defined as the region of interest (ROI) [6, 9]. In general, the average signal of the pixels detected in a given ROI is used as the calcium activation signal of the target neuron. Detecting the pixels of neuronal cell bodies with active calcium signals is straightforward; however, depending on the experimental conditions and size of the target structure, the signal-to-noise power ratio (SNR) of calcium signals is often low.

A reduction in the SNR of calcium imaging is mainly caused by the low expression performance of the calcium indicator, short dwell time of each neuron, small target structure, or a combination of these factors [10, 11]. Heterogeneity in calcium indicator expression causes SNR reduction in neurons with low expression. Some neurons with low calcium indicator expression have a low SNR because the calcium activation signal level itself is relatively low [12]. The shorter dwell time causes the SNR to decrease for each neuron [13]. In addition, experimental factors such as inflammatory reactions and blurring due to regeneration of the dura lower the SNR. Neurons with low SNRs often appear as blurry outlines in the mean images of calcium imaging movies; however, the signal extraction performance of these neurons has not been extensively discussed in previous studies.

Efforts to extract the signals of target neurons from calcium imaging data began with principal component analysis (PCA) and independent component analysis (ICA)-based ROI detection methods [14]. The PCA/ICA technique was developed for the effective separation of signals from the overlapped area of neurons; however, the denoising performance was not sufficient for low-SNR data. Since the development of the PCA/ICA technique, many ROI detection methods have been proposed to improve denoising capability. In particular, once the constrained non-negative matrix factorization (CNMF) method based on calcium dynamics modeling was proposed, this method has been mainly used for calcium imaging analyses [15-18]. CNMF is effective for detecting the shape of many overlapping neurons and simultaneously extracting signals from these neurons. Despite these benefits, the signal extraction performance remains insufficient for neurons with a low SNR. The Suit2p method is a specialized technique for analyzing a large number of neurons; however, it has not been extensively analyzed for the detection and denoising of neurons with a low SNR [19].

In calcium imaging, current ROI detection methods detect morphologically clustered pixels in the shape of neurons; however, signals extracted from these ROIs are susceptible to noise [20, 21]. Signals from pixels that cover intracellular organelles, where there is no calcium indicator, will reduce the SNR, despite the pixels being clustered morphologically. Detecting these noisy pixels as an ROI lowers the SNR of the detected pixel’s average signal. Conversely, detecting pixels outside the shape of a neuron as the ROI can reduce baseline noise when calculating the average of the ROI pixels. Detecting the ROI as a criterion to increase the SNR can improve signal extraction performance.

In this paper, an iterative correlation-based ROI detection (ICoRD) method is proposed, which enhances the SNR of target neurons by iteratively estimating the ground-truth calcium signal (true activation calcium signal without noise). ICoRD detects pixels that maximize the correlation coefficient between the estimated ground-truth calcium signal and the average signal of the detected pixels. The performance of ICoRD was verified on a simulated dataset with the ground-truth calcium signal, and ICoRD was applied to *in vivo* mouse calcium images to validate the signal extraction performance. The ICoRD results were compared with those of CNMF. Compared to CNMF, ICoRD showed substantially higher signal extraction performance, which was evaluated by calculating the correlation coefficient with the ground-truth calcium signal, especially when the SNR was lower than -15 dB. The lower the SNR of the raw data, the greater the difference in performance between the two methods. ICoRD will provide neuroscientists with reliable calcium signal extraction results, even in neurons with a low SNR.

## 2. Method

### 2.1. Overview of the ICoRD for extracting neural signals in calcium imaging data

A novel iterative correlation-based ROI detection method that iteratively detects ROIs to obtain signals closer to the ground-truth calcium signal is proposed (Figure 1(a)). The first reference signal for the iterative update was defined as the average signal of nine pixels adjacent to a user-defined pixel in the neuron of interest. Within the observation range (red dotted box in Figure 1(a)), which is a group of pixels in which the cell body of the neuron can be best visualized within the surrounding space, pixels were selected to maximize the correlation coefficient between the average signal of the selected pixels and the reference signal.

**Figure 1.**
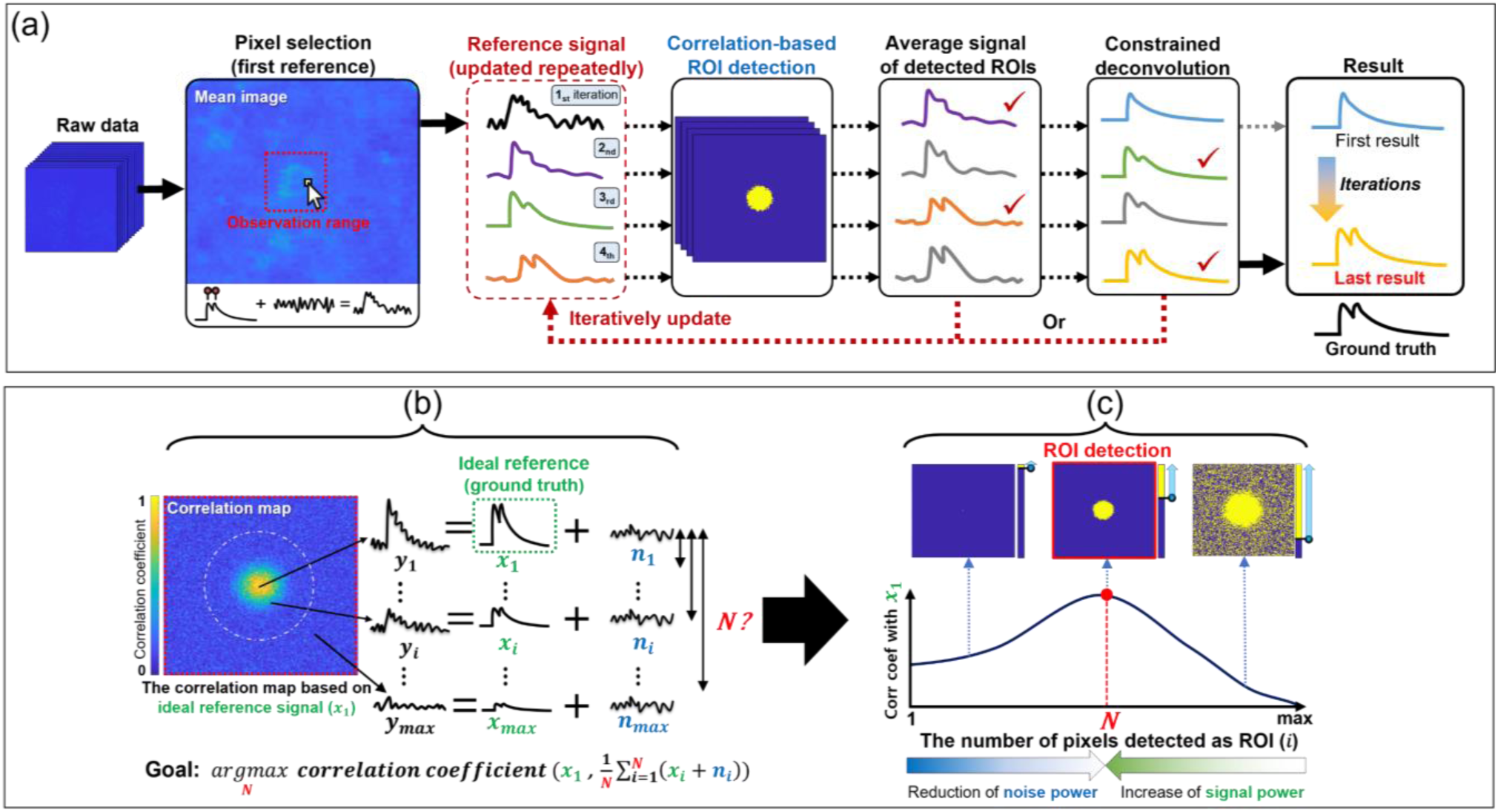
Schematic representation of the proposed ICoRD algorithm for neural signal extraction. (a) Framework of the ICoRD algorithm for estimating accurate activation signals from the target neuron. A representative pixel and observation range were manually selected (left). ICoRD used the average calcium signal from nine pixels adjacent to the selected pixel as the first reference signal for ROI detection. The reference signal was iteratively updated to approximate the ground-truth calcium signal. For ROI detection, a correlation-based ROI detection method ((b)–(c)) was applied, which detected the ROI to have an average signal most similar to the reference signal. The average signal of the pixel corresponding to the ROI and the constrained deconvolution (CD) signal were computed in each iteration. The reference signal of the next iteration was selected from the average signal or CD of the previous iteration. When the iteration was completed, the CD signal of the final iteration was output as the inferred signal of the target neuron (right). (b) Correlation-based ROI detection to obtain the set of pixels that maximizes the correlation coefficient between the average signal of the detected pixels and the reference signal. (c) The optimal number of detected pixels (*N*) of the ROI was determined to maximize the correlation coefficient between the reference signal and the averaged ROI signal.

To detect the ROI that maximizes the correlation coefficient with a reference signal, we calculated the correlation coefficients twice in total. For the first calculation, ICoRD calculated the correlation coefficients between the reference signal and the signals of individual pixels to sort the pixels. All candidate pixels were sorted in descending order of the correlation coefficient calculated based on the reference signal (the number of pixels *i* ranged from one to the maximum number of pixels (*max*) within the observation range in Figure 1(b)). The second correlation coefficient calculation was performed to calculate the correlation coefficient between the reference signal and the average signal of the different number of pixels (*i* = 1 ∼ *max*) in Figure 1(c). The number of pixels that maximizes the correlation coefficient with the reference signal was defined as the optimal number of pixels (*N*), and a pixel group with an optimal number of pixels was detected as an ROI. In the process of obtaining this second correlation coefficient, we observed a trade-off between signal power increase and noise power reduction. When the number of pixels was significantly large, pixels with weak signals were averaged, which reduced the correlation coefficient with the reference signal (Figure 1(c)). However, when the number of pixels was significantly small, white noise was pronounced, resulting in a low correlation coefficient (Figure 1(c)). A set of pixels with the optimal number of pixels was chosen as an ROI to achieve the highest correlation. The ground-truth calcium signal was then estimated by iteratively setting a new reference signal as the average signal of the ROI or the deconvolution signal of the average signal (Figure 1(a)). This process was repeated until the reference signal was close to the ground-truth calcium signal. The detailed iteration algorithm is shown in Figure S2.

### 2.2. Initial parameter setup

To select the first reference signal, the ICoRD algorithm began by selecting the representative pixel of the target neuron. The average signal of the nine pixels adjacent to the selected pixel was used as the first reference signal. The observation range that best visualized the neuron was selected by checking the movie, mean image, and correlation image. The mean image represented the average time-axis signal for each pixel. The correlation image was computed by averaging the correlation coefficient of each pixel with its four immediate pixels. When certain pixels had to be excluded for various reasons, such as overlapping with neighboring neurons, the algorithm was executed without the rejection range selected by the user (Figure S1). The initial parameters used in CNMF are described in the Supporting Information (Table S1).

### 2.3. Iterative update of the reference signal

In ICoRD, the correlation coefficients are calculated between the initial reference signal of the target neuron and all signals in each pixel within the observation range. To detect the ROI that maximizes the correlation coefficient with the ground-truth calcium signal, a correlation-based ROI detection method is proposed (Figure 1(b)–(c)). The correlation-based ROI detection method only detects pixels of an optimal number as an ROI.

Thus far, we have described how to determine the optimal number of pixels that maximizes the correlation coefficient between the average signal of the ROI and the estimated ground-truth calcium signal. To estimate the ground-truth calcium signal of the target neuron *in vivo*, a constrained deconvolution (CD) method was applied to the average signal of the detected ROI [18]. Because this CD method has a denoising effect, its application helped estimate the ground-truth calcium signal. ICoRD iteratively detected the ROI and updated the average signal or CD signal to the reference signal, such that the reference properly approximated the ground-truth signal. The criterion for selecting the reference signal was the information difference between the average calcium signal of the previous and current iterations. The information difference was one minus the correlation coefficient between the average signal of the ROI detected in the current (*Rawc*) and previous iterations (*Rawp*) (Equation 1).

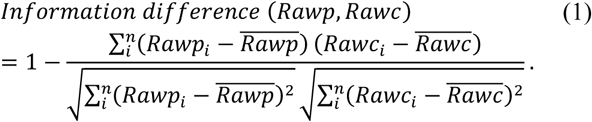

The information difference in each iteration was used to select a reference signal to avoid significant bias in the initial CD result. A detailed algorithm flowchart of ICoRD is presented in Figure S2. ICoRD eventually provided the CD result reconstructed from the average signal of the ROI with the maximum correlation coefficient using the estimated ground-truth calcium signal from the last iteration.

### 2.4. Correlation-based ROI detection

Figure 1(b)–(c) show the correlation-based ROI detection method. This method effectively extracts the calcium signal of a target neuron, even in extremely noisy environments. The ROI detection technique proposed in this study aims to detect a neuron signal closer to the ground-truth calcium signal. ICoRD was assumed to acquire a reference signal close to the ground-truth calcium signal upon iteration, and an ROI detection method that maximized the correlation coefficient with the ground-truth calcium signal was first derived.

In calcium imaging, denoising techniques for individual neurons are evaluated based on the correlation coefficient of the denoised result and the ground-truth calcium signal. However, most existing calcium imaging analysis methods extract neural signals through post-processing after selecting an ROI based on neuron shape. The proposed correlation-based ROI detection method detects the ROI in which the correlation coefficient between the reference signal and the average signal of the ROI is the maximum. In a noise-free environment, the ROI detection method that maximizes the correlation coefficient is used to detect a single pixel as an ROI that maximizes the correlation coefficient of the ground-truth signal.However, in an environment with noise, the ROI detection method that maximizes the correlation coefficient with the ground-truth calcium signal should consider the white Gaussian noise reduction effect by averaging the signals of the pixels detected as the ROI. Therefore, only pixels of an optimal number, at which the correlation coefficient between the average signal of the ROI and the ground-truth signal was maximized, were detected. The correlation coefficient trend between the ground-truth signal and the average signal of the detected ROI as increases of the number of pixels had a concave shape near the maximum correlation coefficient, as shown in Figure 1(c). The concave shape represents the trade-off relationship between the reduction in noise power owing to the averaging effect and the increase in signal power owing to the detection of a highly correlated signal. Figure 1(c) shows the optimal number of pixels (*N*), which determines the ROI that has the maximum correlation coefficient with the ideal reference signal (*x*_1_) of the target neuron. The temporal signal (*y*) of each pixel (*i*) is composed of a signal (*x*) and noise (*n*). Pixel order (*i*) is sorted in descending order based on the Pearson correlation coefficient of each pixel’s signal (*y*_*i*_) and ground-truth signal (*x*_1_). The correlation-based ROI detection method maximizes the correlation coefficient between the average signal of the ROI and ground-truth calcium signal by detecting pixels with an optimal number (*N*).

## 3. Results

### 3.1. Performance of ICoRD in simulated calcium imaging datasets

Having established a new signal-extraction algorithm, its performance at various noise levels was investigated. For a fair measurement of the signal extraction performance of ICoRD, a ground-truth calcium signal and various modeled noise levels were required. Therefore, a realistic model that simulates calcium imaging data with a designated noise level was developed. All simulations and analyses were performed using MATLAB R2020b (MathWorks, Natick, MA, USA). The simulated calcium imaging data were generated by modeling and multiplying the spatial and temporal components of calcium dynamics. The spatial component was modeled as a Gaussian shape of light propagation owing to the calcium indicator at the central location (75,75) in 150 horizontal and 150 vertical spaces (Figure 2(a)). The temporal component for single activation was modeled as a 10-Hz sampling rate based on the rising tau (*τ*_*r*_ = 550 ± 52 ms) and decaying tau (*τ*_*d*_ = 179 ± 23 ms) values of pGP-CMV-GCaMP6s (GCaMP6s) (Figure 2(b), Equation 2) [22].

**Figure 2.**
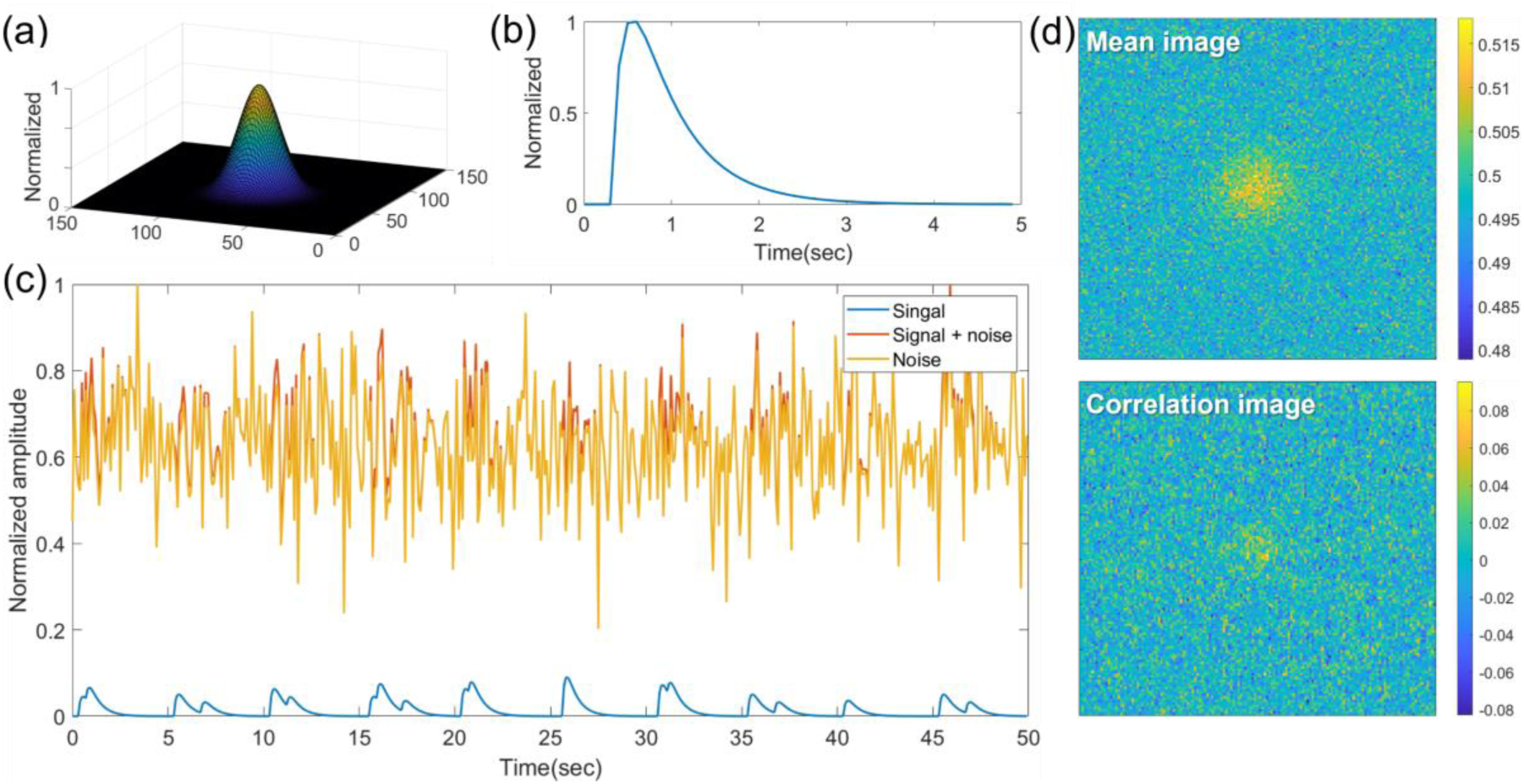
Simulated calcium dynamics at a severe noise level (−30 dB). (a) The spatial component of the calcium dynamics model. (b) The temporal component of the calcium dynamics model derived from the kinetics of GCaMP6s. (c) An example of the simulated calcium trace of the center pixel with an SNR of -30 dB. (d) An example of the mean and correlation images of the simulation data with an SNR of -30 dB.

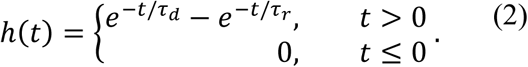

The resulting signal and noise of the central pixel’s simulated calcium trace from the neuron with an SNR of -30 dB are illustrated in Figure 2(c). The ratio of the neuron signal power to the white Gaussian noise power was calculated as the SNR to generate a simulated calcium movie with various SNRs (Equation 3).

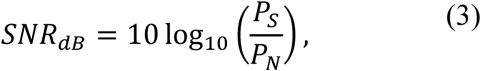

where *P*_*S*_ denotes the signal power, and *P*_*N*_ is the noise power. The modeled temporal dynamics were generated as simulation data for 50 s based on a 10-Hz sampling rate by convoluting the calcium events at various intervals. If selective ROI detection using ICoRD increases the sensitivity of event detection, the events could be resolved in severely noisy images, and closely interleaved events could be resolved better than with the conventional CNMF method. To build a testing platform, two or more events occurring within 5 s with various intervals and noise levels between -30 and 0 dB SNRs were modeled. Figure 2(d) shows the mean and correlation images of the simulated calcium movie with an SNR of -30 dB. In general, the mean image provides an estimate of the location and shape of the neuron because the average pixel value of the neuron signal tends to be higher than the background signal. In the correlation image, the more frequent the activation of neurons or the stronger the signal intensity versus noise, the clearer the appearance of the neuron.

The performance of the proposed ICoRD algorithm was compared with that of CNMF using the above-mentioned simulated calcium imaging data. A comparison was conducted between the extracted traces of the ROIs set by the two methods from the simulated calcium images with an SNR of -30 to 0 dB (Figure 3). In the CNMF method, if the selected ROI candidate was determined as unsuitable, it was automatically removed. Using the calcium images with severe noise (−30 dB), CNMF automatically determined and rejected all the pixels in the image with an SNR of -30 dB as noise. Therefore, we restored the ROIs by recovering the candidate ROIs that were originally rejected by CNMF for comparison with ICoRD. Conversely, ICoRD detected most pixels inside a neuron, although a substantial number of background pixels were included within the ROIs (Figure 3(a)). Furthermore, the extracted signal from the ICoRD-selected ROI correlated significantly better with the ground-truth than the detected ROI from CNMF (Figure 3(b); the correlation coefficients with the ground-truth from CNMF and ICoRD were 0.8267 and 0.9834, respectively). Subsequently, the performance of ICoRD at various noise levels was tested. At all noise levels, signals extracted from ROIs detected by ICoRD were closer to the ground-truth calcium signal than those extracted by CNMF (Figure 3(c)–(d)), demonstrating that ICoRD-based ROI detection is more robust and suitable for noisy signals, as in many *in vivo* calcium images. As the neuron’s SNR decreased, ICoRD lowered the noise level by detecting not only the pixel corresponding to the neuron, but also the surrounding pixels to maximize the correlation coefficient with the reference signal. In the case of CNMF, all ROIs in the data with SNRs of -30 dB and -25 dB were automatically determined as noise and rejected; therefore, the rejected ROIs were restored. CNMF detected a smaller number of pixels as the SNR decreased. Figure 3(d) shows the correlation coefficient of the ground-truth signal and the signal extracted by the two methods from 20 datasets (140 total) with white Gaussian noise added to each SNR (−30 to 0 dB). At all noise levels, the ICoRD result had a higher correlation coefficient with the ground-truth signal than the CNMF result. As shown in Figure 3(d), the difference in performance between the two methods was particularly large in the SNR range of -30 dB to -20 dB. Because the proposed method is a correlation-based ROI detection method that aims at true calcium signal extraction, it shows excellent performance at severe noise levels (SNRs of -30 to -20 dB).

**Figure 3.**
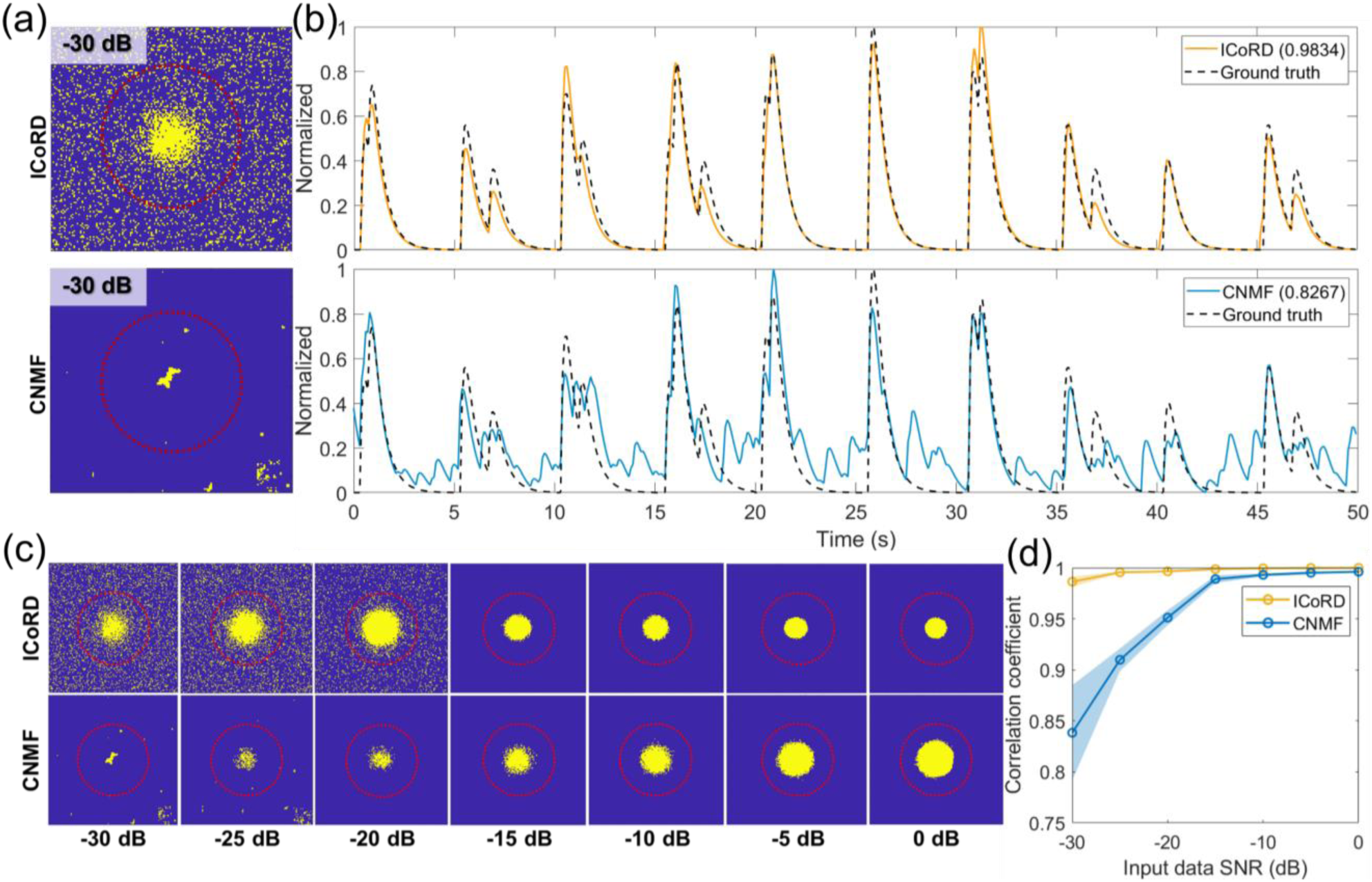
Results of the proposed ICoRD method and CNMF in simulated calcium imaging datasets. (a) The pixels included in the ROIs from the simulated calcium images (yellow) with an SNR of -30 dB using ICoRD (upper) and CNMF (lower). The red dotted line indicates the range in which the neuron’s signal exists in a Gaussian shape. (b) The extracted calcium traces from the selected pixels of the simulated calcium images with an SNR of -30 dB using ICoRD (upper) and CNMF (lower). (c) Detected ROIs using ICoRD (upper) and CNMF (lower) from the simulated calcium imaging data with SNRs of -30 to 0 dB. The red dotted lines are as in (a). (d) Mean (solid line) and standard deviation (shading) of the correlation coefficient between the ground-truth and extracted signals using ICoRD (yellow) and CNMF (blue).

### 3.2. Parameters that change during iteration in ICoRD

Figure 4 shows the trend of the estimated and ground-truth correlation coefficients and the information difference during the iteration of the proposed method in simulated calcium imaging data with an SNR of -30 dB. The estimated correlation coefficient at each iteration represents the correlation coefficient between the constrained deconvolution (CD) result of the last iteration and the CD result of each iteration. The ground-truth correlation coefficient of each iteration is the correlation coefficient between the ground-truth signal and the CD result of each iteration. ICoRD uses the average signal (Avg) of the ROI or CD result as a reference signal for the next iteration. These two reference sources determine the calcium signal extraction performance of ICoRD. Figure 4 shows the estimation performance of the ground-truth calcium signal based on reference signal selection. The Avg-based update has a white background, and the CD-based update of the reference has a pink background. Figure 4(a) shows the trend of the estimated and ground-truth correlation coefficients when the reference signal was selected using ICoRD. The trends of the estimated and ground-truth correlation coefficients were similar, and the ground-truth correlation coefficient generally increased with iterations (Figure 4(a)). This trend of the ground-truth correlation coefficient reveals that ICoRD has been effectively designed to estimate the ground-truth calcium signal of the target neuron during the iterations. The maximum ground-truth correlation coefficient was 0.9889 at the 17^th^ iteration. The bottom of Figure 4(a) shows the vibration trend of the information difference, which is the criterion for selecting the Avg or CD as the reference signal (Equation 1, Figure S2). The information difference is used for avoiding significant bias in the calcium model estimated by only CD during the initial iteration. If the information difference is less than a certain threshold (for example, 0.02), the algorithm determines that there is no more information to be obtained from the previous reference source and updates it with another data source (Raw→CD, CD→Raw). In the process of information difference oscillation, ICoRD estimates closer to the ground-truth calcium signal. An Avg-based update could estimate the ground-truth signal with a moderately high correlation coefficient (0.9597 at the 17^th^ iteration); however, the estimation performance was relatively low compared to ICoRD owing to the lack of the noise removal effect in CD (Figure 4(b)). In addition, in the case of Avg-based update, the improvement of the ground-truth correlation coefficient by iteration was hardly observed after 6^th^ iteration. Although CD has a noise-removing effect, there is a risk of continuously estimating the calcium model at the initial iterations in CD-based updates (Figure 4(c)). In the CD-based update, the speed of finding the ground-truth calcium signal as increases of the iteration was relatively slow, and the maximum ground-truth correlation coefficient was 0.8087 in the 14^th^ iteration, which was not sufficient to analyze neural activity. The information difference of the CD-based updates was saturated to zero overall with iterations. This saturated information difference indicates that the calcium model incorrectly estimated in the initial iteration has been maintained continuously from the initial iteration. The results in Figure 4 show that appropriate conversion of the reference source allows ICoRD to estimate the ground-truth signal with a high correlation coefficient.

**Figure 4.**
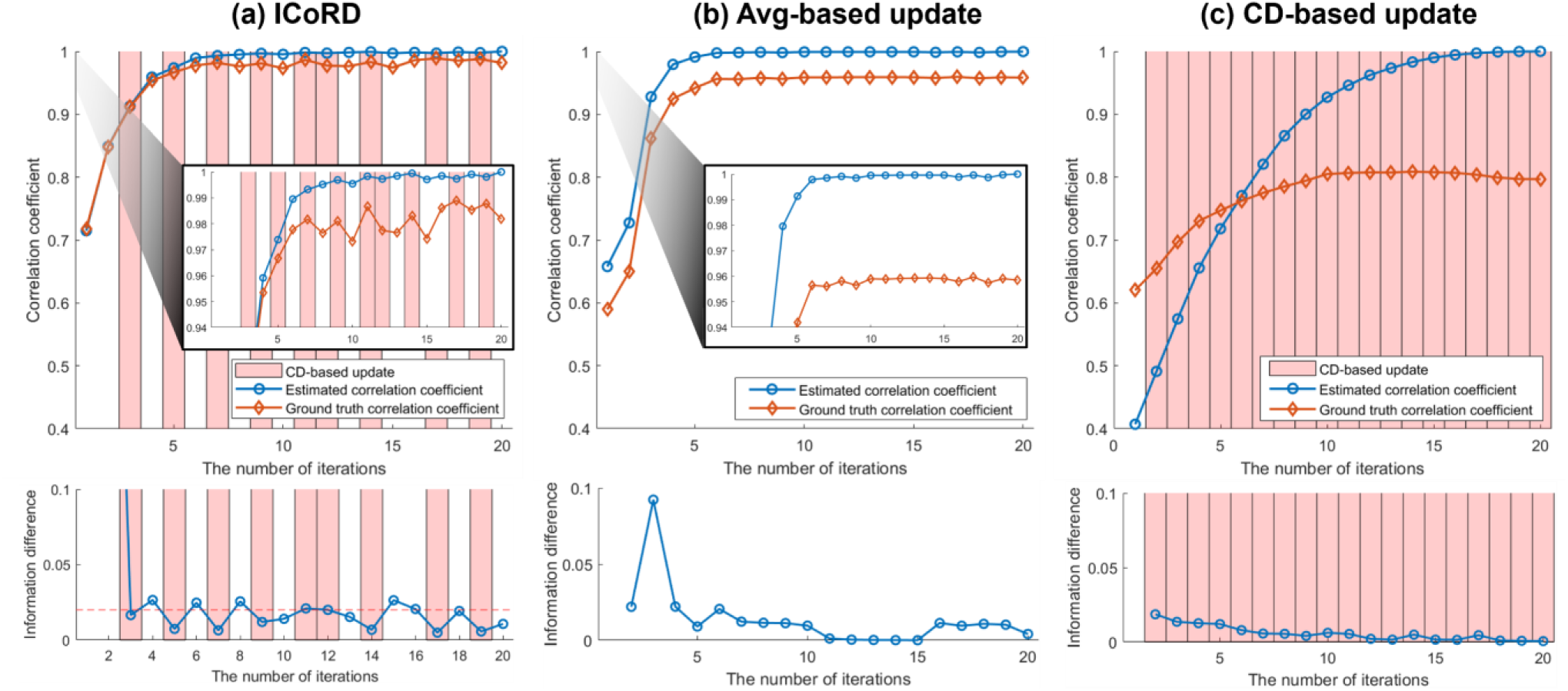
Trend of estimated correlation coefficient, ground-truth correlation coefficient, and information difference during iterations in the simulation data with an SNR of -30 dB. (a) Comparison of estimated and ground-truth correlation coefficients during iterative reference updates in ICoRD. (b) Comparison of estimated and ground-truth correlation coefficients during iterative reference updates based on the average (avg) signal of the detected ROI. (c) Comparison of estimated and ground-truth correlation coefficients during iterative references updates based on CD results.

### 3.3. Application of the ICoRD algorithm to in vivo two-photon calcium imaging data

It was verified that the proposed method has a powerful advantage in terms of the SNR in the simulation data. In this section, the practicality of the proposed method is examined using two-photon calcium imaging data measured in a mouse brain.

All procedures involving mice were approved by the Korea Brain Research Institute, Institutional Animal Care and Use Committee (approval number: IACUC-19-00041). Adult (older than P60) transgenic mice expressing GCaMP6s (C57BL/6J-Tg [Thy1-GCaMP6s] GP4.12Dkim/J, The Jackson Laboratory) underwent surgery to implant a cranial window and headplate. A 3-mm craniotomy was performed over the posterior parietal cortex (PPC) area centered 2 mm posterior and 1.7 mm lateral to the bregma. The window covering the craniotomy area was constructed by bonding 3-mm and 5-mm diameter No. 0 coverslips (Warner Instruments). The titanium headplate was attached to the skull over the window using opaque dental cement (Super-Bond, Sun Medical) and covered all exposed tissue.

Images were acquired with a Nikon 16X (0.8 NA with 3.0 mm working distance) objective lens attached to a microscope (HyperScope, Scientifica) running ScanImage (2019a). A Ti-Sapphire LASER (Chameleon Vision II, Coherent) operated at 910 nm was used as the light source for the imaging microscope. The head-fixed mouse was awake during the imaging procedure of the PPC and kept warm with a custom-made body restrainer. The imaging area was approximately 200 μm deep and centered at the PPC; it was adjusted to avoid thick blood vessels appearing in the ROI. The resolution of the images was 512 × 512 pixels, and the FOV was a 700 μm square. The frame rate of the microscope was set to 30-Hz.

The proposed ICoRD algorithm was applied to raw *in vivo* calcium imaging data using the same algorithmic process as the simulation-based validation. The spatial and temporal results of the selected neurons using ICoRD and CNMF *in vivo* calcium imaging data were compared (Figure 5). To compare ICoRD with CNMF, a neuron with high and frequent calcium activity (neuron 1) and a neuron with relatively sparse activity (neuron 2) were selected among the neurons clearly detected by CNMF (Figure 5(a)). A neuron with a low SNR and sparse activation (neuron 3) was detected by ICoRD but not CNMF. Figure 5(b) shows the zoomed images (mean, correlation) around the three target neurons in the *in vivo* calcium imaging movie and the ROIs detected by ICoRD and CNMF. The location of the target neuron is marked in the mean and correlation images by a red dotted line. The SNR of neuron 1 was sufficiently high to be identified in both the mean and correlation images. The ROI of neuron 1 detected by ICoRD and CNMF were similar. The results of the two methods for high-SNR neurons *in vivo* were similar to the detection trends observed in the simulation environment. Neuron 2 only exhibited the blurry shape of the target neuron in the mean image. ICoRD detected clustered pixels and several surrounding pixels outside the target neuron, whereas CNMF detected only the clustered pixels of neuron 2. Neuron 3 only showed a blurry neuron morphology in the mean image and its ROI was only detected by ICoRD. ICoRD extracted signals from neuron 3 despite this neuron being sparsely activated at high noise levels. ICoRD detected the ROI of neuron 3 with clusters and surrounding pixels, similar to the ROI of neuron 2. These ROI detection results with the pixels surrounding neurons 2 and 3 were similar to the ROI detection results in the simulation verification with low SNRs. Because ICoRD detects the ROI that maximizes the correlation coefficient with the estimated reference by detecting pixels that have a high correlation with the estimated reference signal, it detects surrounding pixels in the case of very low SNR neurons and takes advantage of the noise reduction effect.

**Figure 5.**
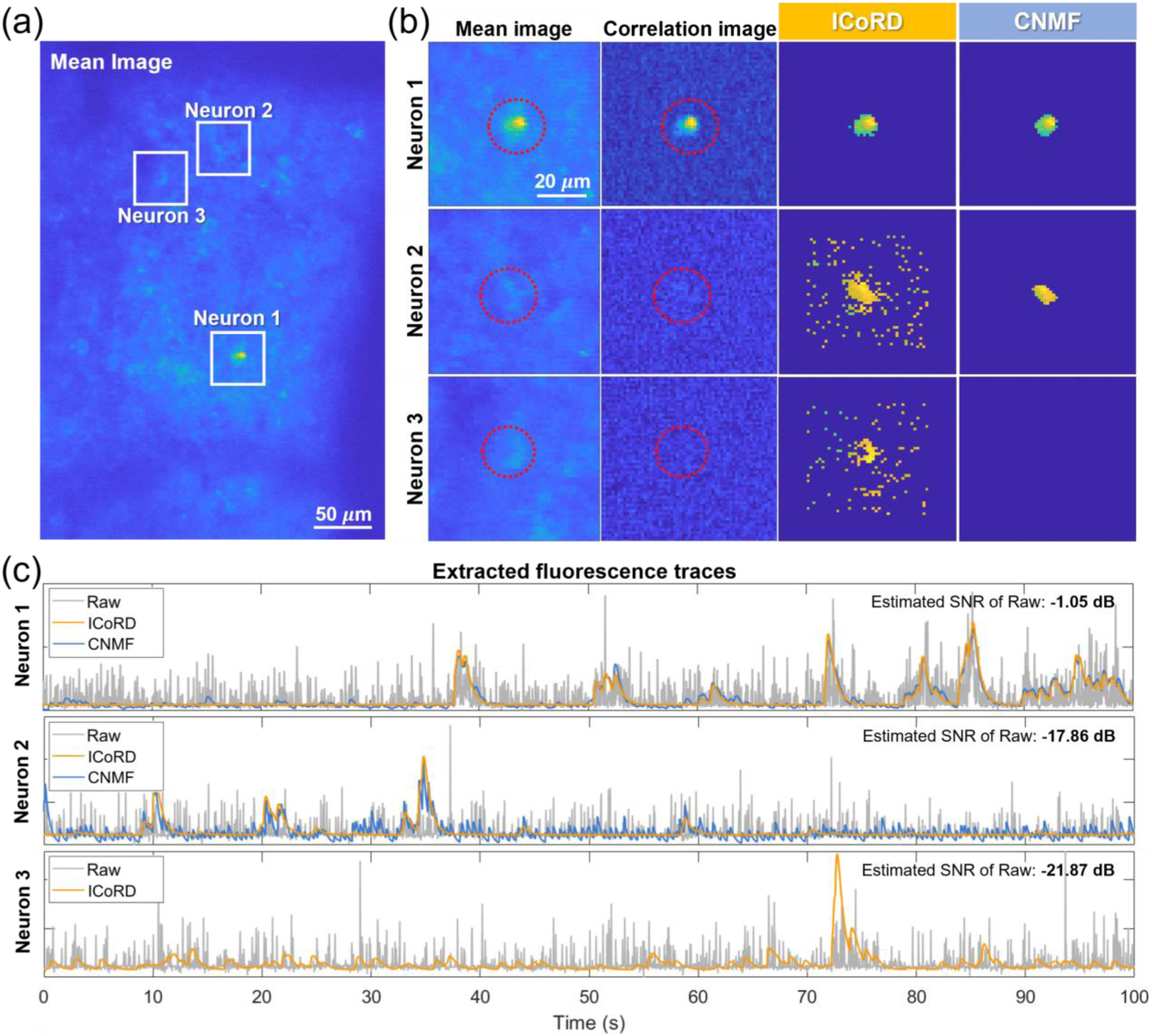
Application to *in vivo* two-photon calcium imaging data. (a) Mean image of the *in vivo* calcium imaging movie and three target neurons. (b) The zoomed mean and correlation images around the target neurons (1,2,3) and the ROIs detected by ICoRD and CNMF. (c) The first reference (average signal of 9 pixels adjacent of the selected pixel) of the target neurons (1,2,3) and extracted temporal result using ICoRD and CNMF. The estimated SNR of neuron 1, neuron 2, and neuron 3 were -1.05 dB, -17.86 dB, and -21.87 dB, respectively.

Figure 5(c) shows the raw signal of the selected pixel as a ‘Raw’ (grey line) of the three neurons in the ICoRD algorithm and each CD result of the ROI detected by ICoRD (orange line) and CNMF (blue line). We present the first reference source (Raw) and the result of the last iteration of ICoRD to explain how the estimated reference changes with iteration. Each signal was normalized by subtracting the mean and dividing it by the standard deviation. A CD-based SNR estimation method was used to evaluate the neurons in an *in vivo* environment (Figure S3). The estimated SNR of individual neurons in the Raw signal was -1.05 dB in neuron 1, -17.86 dB in neuron 2, and -21.87 dB in neuron 3. The signals extracted by ICoRD and CNMF in neuron 1, with a high SNR and frequent calcium activity, exhibited similar trends (correlation coefficient: 0.9608). However, for neuron 2, with a relatively infrequent and low-amplitude calcium activation signal, the signal trends detected by the two methods were relatively different (correlation coefficient: 0.7241). Furthermore, only ICoRD, not CNMF, detected neuron 3 as an ROI. This result is similar to that of CNMF, indicating low signal extraction performance for an SNR below -20 dB in a simulation environment. Although the calcium signal of neuron 3 was highly sparse at high noise levels, ICoRD effectively extracted the calcium trace after iteration. Although ICoRD and CNMF apply the same CD method, the signal obtained from ICoRD appeared to have a higher SNR. This is because ICoRD provides a better input signal for the CD method by detecting the ROI that maximizes the correlation coefficient with the predicted ground-truth signal of the target neuron. ICoRD was confirmed to effectively denoise signals from neurons with a low SNR in *in vivo* calcium imaging data.

## 4. Discussion

In the field of neuroscience, the calcium imaging technique is actively used because the activities of neurons can be assessed simultaneously in a target region. When attempting to observe neurons, those with low SNRs are inevitably observed; however, analyzing these low-SNR neurons remains a challenging task. Existing ROI detection methods for calcium imaging data are aimed at the automatic detection of many neurons; hence, the extraction of signals from neurons with low SNRs often fails. If the calcium signals of neurons with sparse and weak activity can be extracted, the results of various calcium imaging experiments can be analyzed more reliably using the signals of more neurons. Therefore, we developed ICoRD to reliably estimate ground-truth calcium activity, even in neurons with a low SNR.

The proposed ICoRD method effectively extracts calcium signals, even from calcium imaging data with a low SNR. ICoRD performed reliable signal extraction (correlation coefficient with ground-truth calcium signal > 0.98) of the target neuron from simulated calcium imaging data with an SNR of -30 dB to 0 dB, which was difficult to analyze using the conventional method. Because ICoRD successfully estimated the ground-truth calcium signal as the number of iterations increases, it was possible to detect the ROI that enhanced the ground-truth correlation coefficient based on the estimated reference. ICoRD was confirmed to robustly extract activation signals, even from neurons with sparse activity in an *in vivo* environment with severe noise levels.

Because the proposed algorithm aims to extract temporal calcium activity, the focus of this study was not on detecting the shape of the spatial component. The proposed method clearly estimated the signal of a neuron with a low SNR, which was difficult to achieve using the conventional method. ICoRD detected neurons with very sparse activity, which were blurry in the mean image and not visible in the correlation image. The robust extraction performance of ICoRD may have been possible because the proposed algorithm does not require the ROI to be clustered, like a neuron. The ROI detection method that maximizes the correlation coefficient with the reference signal, regardless of the shape of the ROI, is very effective when extracting the calcium signal from a single neuron.

ICoRD provides calcium signals with a high SNR from the target neuron; however, the user must determine the location of the target neuron for the first reference signal. If the target neuron significantly overlaps with other neurons, the user must choose a range of pixels to reject. These initial parameters of ICoRD can be laborious if thousands of neurons must be detected. Therefore, ICoRD is recommended when users need to extract signals from hundreds or fewer individual neurons with low SNRs. If the user identifies the target neuron in two-photon calcium imaging data but the conventional method fails to detect neurons with low SNRs, ICoRD provides a calcium trace with high reliability.

## 5. Conclusion

ICoRD repeatedly estimates a reference close to the ground-truth calcium signal and obtains a pixel group that maximizes the correlation coefficient with the reference signal. Because ICoRD aims to extract the calcium signal that maximizes the correlation coefficient with the ground-truth calcium signal of the target neuron, regardless of the shape of the ROIs, it extracts calcium signals from neurons with a low SNR with a high level of reliability compared with the existing technique. The signal extraction performance of ICoRD in simulated neurons with various SNR was verified, and its applicability was tested *in vivo*. The ROI detection idea that maximizes the correlation of ICoRD provides a better calcium signal than existing calcium signal-based spike deconvolution studies. By analyzing neurons with a low SNR via ICoRD, we expect to better understand the circuit-wide neural dynamics relevant to the function of the brain.

## Supporting Information for

### Demixing of overlapped neurons using iterative correlation-based region of interest (ROI) detection (ICoRD)

When the target neuron overlaps other neurons, ICoRD executes the algorithm after excluding the overlapping pixels. If the user can identify two overlapping neurons, ICoRD can distinguish their signals. Figure S1A shows that the rejection range was determined when overlapping neurons (neuron A: left neuron, neuron B: right neuron) were found in the compressed image (mean, max). Figure S1B shows the result of selecting the ROI of neuron A among the pixels, excluding the rejection range. Figure S1C shows the 100-s ground-truth signals of the two simulated neurons. Figure S1D shows the result of detecting neuron A through ICoRD and the ground truth. ICoRD was able to extract the signal of the desired neuron between the two overlapping neurons in a noisy environment (−10 dB) with high accuracy (correlation coefficient with ground truth: 0.9884).

### Details of the ICoRD algorithm

For each iteration, the reference signal was updated with the average signal (Avg) of the detected ROI or the constrained deconvolution (CD) result in the previous iteration (Figure S2). For the initial iteration, the Avg signal of the ROI was updated with the reference signal because the estimation accuracy of CD in the first iteration tended to be low. However, repetitive updates to only the Avg signal can easily reach a local optimum because it does not utilize the noise-reducing effect or estimating function of CD. Iterative reference updates based only on the CD result can repeatedly find the ROI biased toward the initially estimated calcium dynamics model. Thus, starting from the second iteration, the Avg signal or CD result was selectively updated as a reference signal according to the information difference between each temporal Avg result of the current and previous iterations (Equation 1).

The information difference was used for avoiding significant bias in the calcium model estimated by CD during the initial iteration. When the information difference is less than a certain threshold (for example 0.02), the algorithm determines that no more information is available from the Avg-based iteration and updates the CD result in the next iteration. The CD-based update also changes to an Avg-based update in the next iteration when the information difference falls below a threshold. The estimated correlation coefficient at each iteration represents the correlation coefficient between the CD result of the last iteration and that of each iteration. The algorithm stops at a predetermined maximum number of iterations. At this point, the CD result is output owing to the algorithm.

### Signal-to-noise power ratio (SNR) estimation of calcium imaging data

To estimate the SNR of calcium activity (Raw), the result of CD was defined as a signal, and the data of the lower 25% of the temporal raw data subtracted by CD were defined as noise. We referred to a previous study using the lower 25% amplitude to estimate the noise level when the signal remained in the raw calcium data [1]. Equation S1 represents the proposed SNR estimation equation.

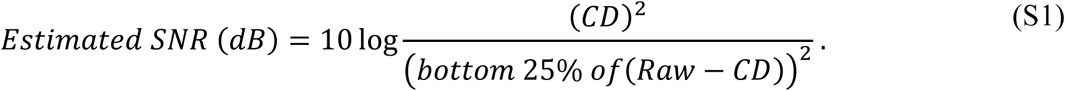

The SNR estimation performance was verified by generating simulation data every 0.2 dB from -30 to 10 dB. All the simulation data generation conditions were the same as those shown in Figure 2. In Figure S3, defining the bottom 25% of Raw-CD as noise revealed a better SNR estimation performance than when defining Raw-CD as noise. The total estimation error of the proposed method at -30 dB to 10 dB was 1.3493 ± 2.2625 dB. Because CD cannot perfectly estimate the signal, the signal may remain after subtracting CD. Therefore, the noise power was greater because part of the signal was determined as noise, and the larger noise power resulted in a lower estimation of the SNR. Using the SNR estimation method, we compared the relative SNRs of the calcium signals of target neurons within the *in vivo* calcium imaging data.

### Constrained non-negative matrix factorization (CNMF) parameters

The CNMF algorithm was applied in this study using the CAIMAN open-source calcium imaging toolbox. The parameters required for the CNMF algorithm were selected according to the guidelines and verified the researchers [2]. The main CNMF parameters used in this study are as follows:

**Table S1.** CNMF parameters.

|  |  |
| --- | --- |
| Sampling frequency (Hz) | 10 (for in silico), 30 (for in vivo) |
| Rising & decaying tau (ms) | 0.179, 0.550 |
| Order of the autoregressive model | 2 |
| Patch size (pixel, pixel) | (32, 32) |
| Merging threshold | 0.8 |
| Neuron size | 5 |

The sampling frequency was based on the values obtained during the simulation and *in vivo* data measurement, and the time constants refer to the results of the GCaMP 6s rodent experiments [3]. To perform deconvolution, the order of the autoregressive model was set to two. The patch size, merging threshold, and neuron size were defined by researchers through parameter optimization to be appropriate for the data.

**Figure S1.**
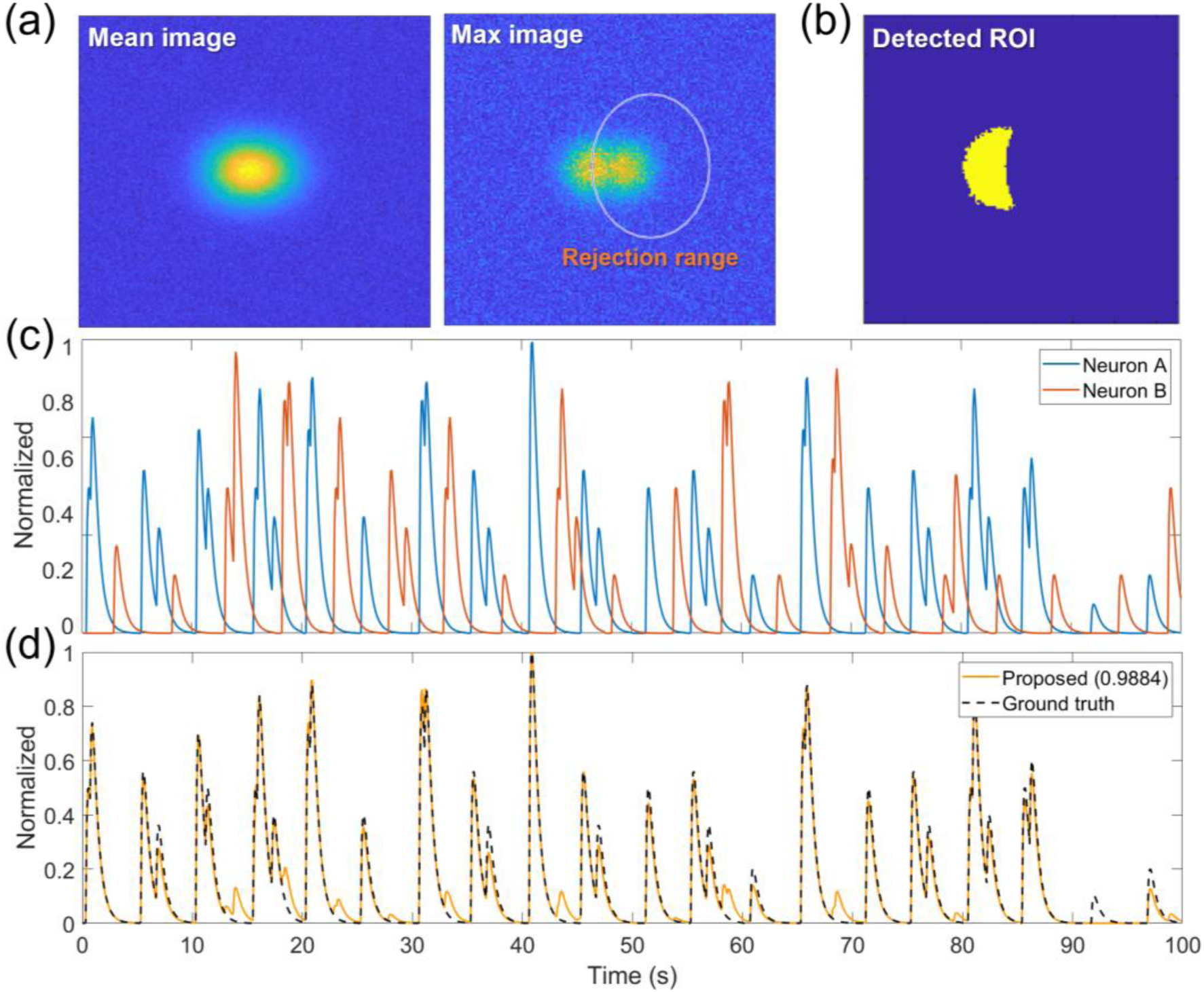
Demixing of two overlapped neurons (Neuron A, Neuron B) in simulation data using ICoRD. (a) Mean and max images of two overlapped neurons with an SNR of –10 dB. (b) Detected ROI of neuron A using the ICoRD method. (c) Simulated temporal calcium dynamics of the two neurons (Neuron A, Neuron B). (d) Temporal result of the detected ROI of neuron A using the ICoRD algorithm.

**Figure S2.**
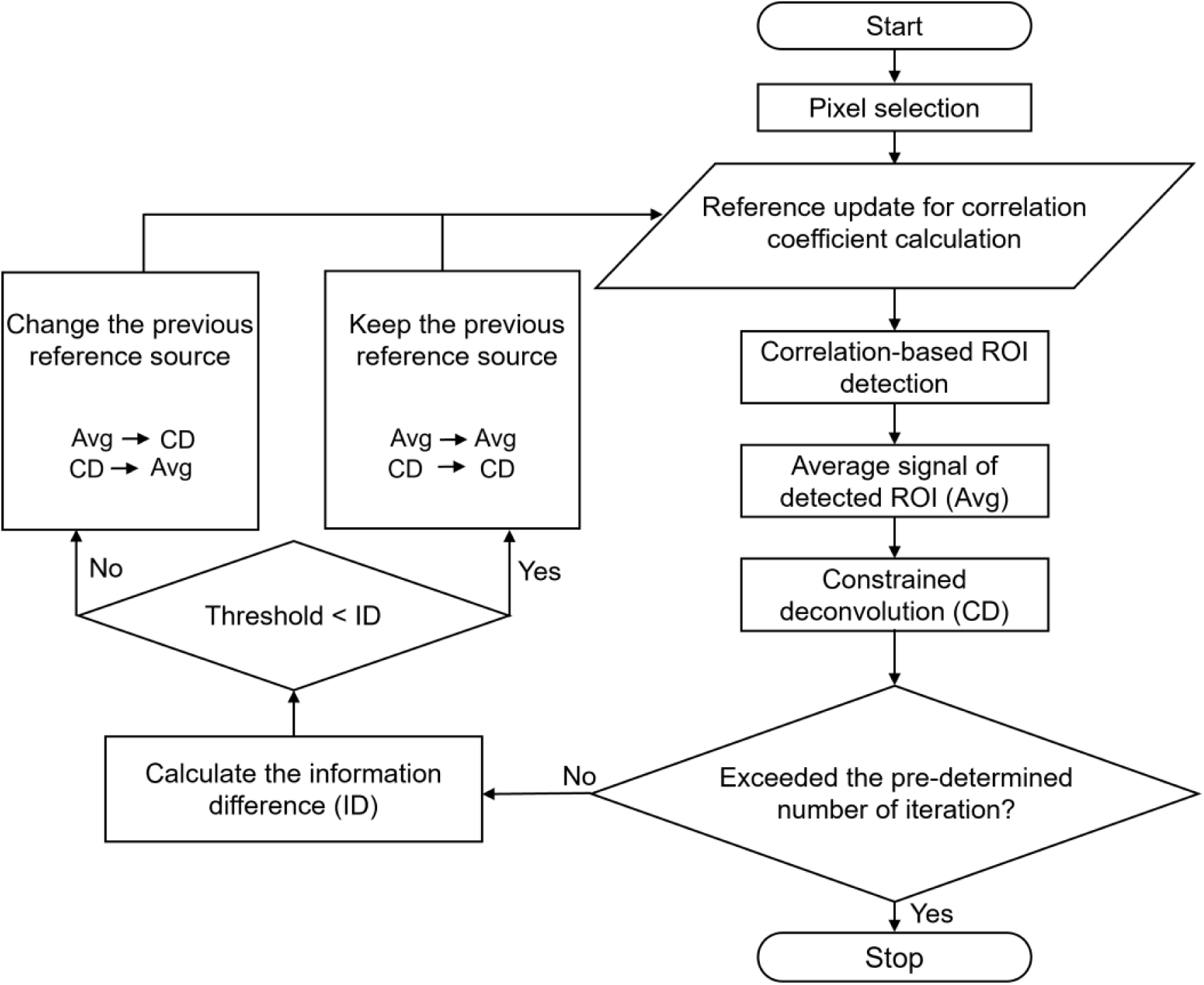
Detailed flowchart of the proposed ICoRD algorithm.

**Figure S3.**
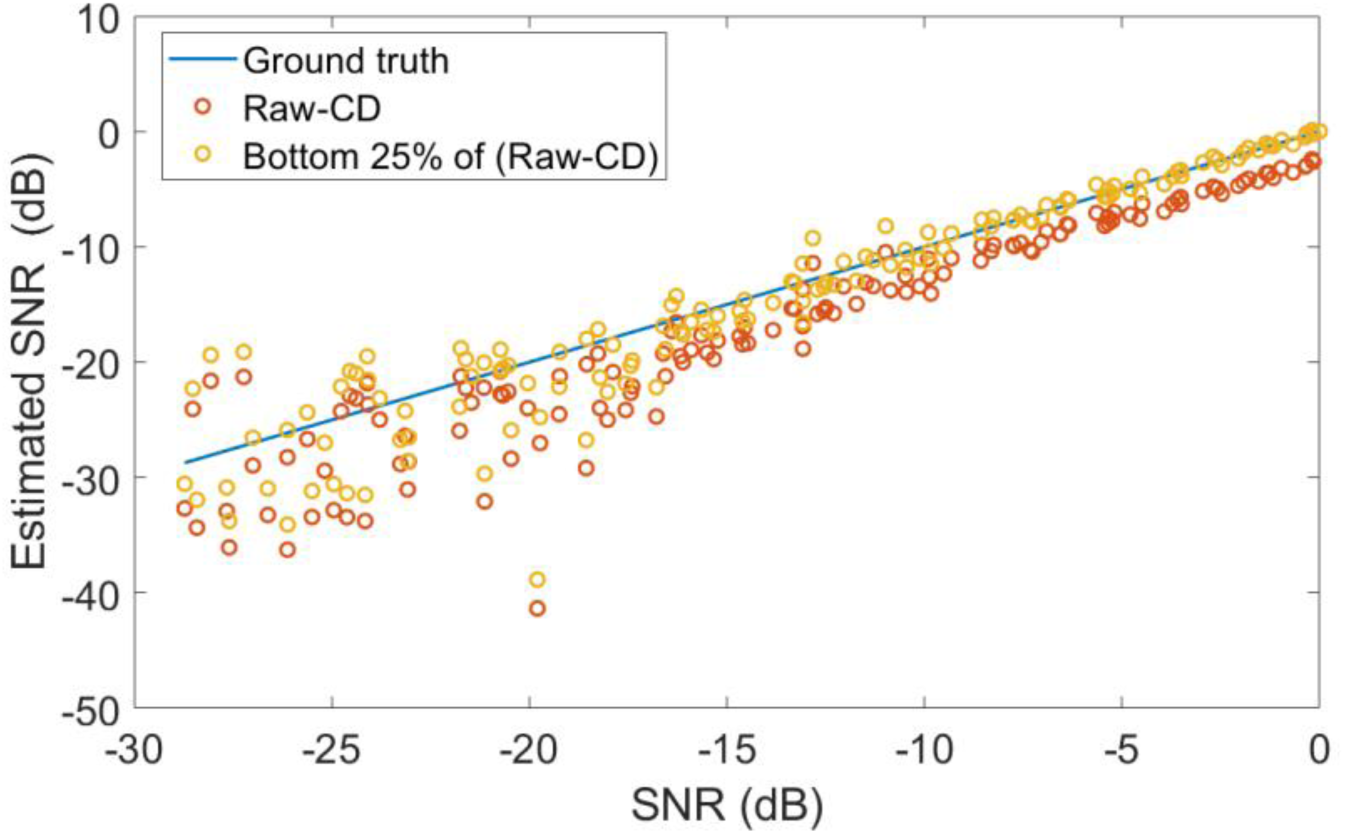
Comparison of estimated SNR with different SNR estimation methods and the ground-truth SNR. The blue line indicates the ground-truth SNR of the simulated calcium traces. The red circles represent the SNR estimation results when the Raw-CD is defined as the noise for each simulated calcium trace. The yellow circles show the SNR estimation results when the lower 25% of the Raw-CD is defined as noise for each simulated calcium trace.

## References

1. Clapham DEJC. Calcium signaling. Cell. 2007;131(6):1047–58.

2. Berridge MJ, Bootman MD, Roderick HLJNrMcb. Calcium signalling: dynamics, homeostasis and remodelling. 2003;4(7):517–29.

3. Helmchen F, Imoto K, Sakmann B. Ca2+ buffering and action potential-evoked Ca2+ signaling in dendrites of pyramidal neurons. Biophys J. 1996;70(2):1069–81.

4. Maravall M, Mainen ZF, Sabatini BL, Svoboda K. Estimating intracellular calcium concentrations and buffering without wavelength ratioing. Biophys J. 2000;78(5):2655–67.

5. Tsien RYJN. A non-disruptive technique for loading calcium buffers and indicators into cells. 1981;290(5806):527–8.

6. Grienberger C, Konnerth AJN. Imaging calcium in neurons. 2012;73(5):862–85.

7. Knopfel T. Genetically encoded optical indicators for the analysis of neuronal circuits. Nat Rev Neurosci. 2012;13(10):687–700.

8. Grienberger C, Konnerth A. Imaging Calcium in Neurons. Neuron. 2012;73(5):862–85.

9. Gobel W, Helmchen FJP. In vivo calcium imaging of neural network function. 2007;22(6):358–65.

10. Harris KD, Quiroga RQ, Freeman J, Smith SL. Improving data quality in neuronal population recordings. Nat Neurosci. 2016;19(9):1165–74.

11. Stringer C, Pachitariu M. Computational processing of neural recordings from calcium imaging data. Curr Opin Neurobiol. 2019;55:22–31.

12. Tian L, Hires SA, Mao T, Huber D, Chiappe ME, Chalasani SH, et al. Imaging neural activity in worms, flies and mice with improved GCaMP calcium indicators. Nat Methods. 2009;6(12):875–U113.

13. Yang W, Yuste RJNm. In vivo imaging of neural activity. 2017;14(4):349–59.

14. Mukamel EA, Nimmerjahn A, Schnitzer MJ. Automated analysis of cellular signals from large-scale calcium imaging data. Neuron. 2009;63(6):747–60.

15. Giovannucci A, Friedrich J, Kaufman M, Churchland A, Chklovskii D, Paninski L, et al. OnACID: Online analysis of calcium imaging data in real time. 2017;30.

16. Zhou P, Resendez SL, Rodriguez-Romaguera J, Jimenez JC, Neufeld SQ, Giovannucci A, et al. Efficient and accurate extraction of in vivo calcium signals from microendoscopic video data. 2018;7:e28728.

17. Maruyama R, Maeda K, Moroda H, Kato I, Inoue M, Miyakawa H, et al. Detecting cells using non-negative matrix factorization on calcium imaging data. Neural Networks. 2014;55:11–9.

18. Pnevmatikakis EA, Soudry D, Gao Y, Machado TA, Merel J, Pfau D, et al. Simultaneous Denoising, Deconvolution, and Demixing of Calcium Imaging Data. Neuron. 2016;89(2):285–99.

19. Pachitariu M, Stringer C, Dipoppa M, Schröder S, Rossi LF, Dalgleish H, et al. Suite2p: beyond 10,000 neurons with standard two-photon microscopy. 2017.

20. Klibisz A, Rose D, Eicholtz M, Blundon J, Zakharenko S. Fast, simple calcium imaging segmentation with fully convolutional networks. Deep Learning in Medical Image Analysis and Multimodal Learning for Clinical Decision Support: Springer; 2017. p. 285–93.

21. Apthorpe N, Riordan A, Aguilar R, Homann J, Gu Y, Tank D, et al. Automatic neuron detection in calcium imaging data using convolutional networks. 2016;29.

22. Chen TW, Wardill TJ, Sun Y, Pulver SR, Renninger SL, Baohan A, et al. Ultrasensitive fluorescent proteins for imaging neuronal activity. Nature. 2013;499(7458):295–300.

## References

1. Brondi M, Moroni M, Vecchia D, Molano-Mazon M, Panzeri S, Fellin T. High-Accuracy Detection of Neuronal Ensemble Activity in Two-Photon Functional Microscopy Using Smart Line Scanning. Cell Rep. 2020;30(8):2567–80 e6.

2. Giovannucci A, Friedrich J, Gunn P, Kalfon J, Brown BL, Koay SA, et al. CaImAn an open source tool for scalable calcium imaging data analysis. Elife. 2019;8.

3. Chen T-W, Wardill TJ, Sun Y, Pulver SR, Renninger SL, Baohan A, et al. Ultrasensitive fluorescent proteins for imaging neuronal activity. Nature. 2013;499(7458):295–300.

